# Preservation of eye movements in Parkinson’s disease is stimulus and task specific

**DOI:** 10.1101/2021.08.17.456700

**Authors:** Jolande Fooken, Pooja Patel, Christina B. Jones, Martin J. McKeown, Miriam Spering

## Abstract

Parkinson’s disease (PD) is a neurodegenerative disease that includes motor impairments such as tremor, bradykinesia, and postural instability. Although eye movement deficits are commonly found in saccade and pursuit tasks, preservation of oculomotor function has also been reported. Here we investigate specific task and stimulus conditions under which oculomotor function in PD is preserved. Sixteen PD patients and eighteen healthy, age-matched controls completed a battery of movement tasks that included stationary or moving targets eliciting reactive or deliberate eye movements: pro-saccades, anti-saccades, visually-guided pursuit, and rapid go/no-go manual interception. Compared to controls, patients demonstrated systematic impairments in tasks with stationary targets: pro-saccades were hypometric and anti-saccades were incorrectly initiated toward the cued target in about 35% of trials compared to 14% errors in controls. In patients, task errors were linked to short latency saccades, indicating abnormalities in inhibitory control. However, patients’ eye movements in response to dynamic targets were relatively preserved. PD patients were able to track and predict a disappearing moving target and make quick go/no-go decisions as accurately as controls. Patients’ interceptive hand movements were slower on average but initiated earlier, indicating adaptive processes to compensate for motor slowing. We conclude that PD patients demonstrate stimulus- and task-dependency of oculomotor impairments and propose that preservation of eye and hand movement function in PD is linked to a separate functional pathway through the SC-brainstem loop that bypasses the fronto-basal ganglia network.

**Significance Statement:** Eye movements are a promising clinical tool to aid in the diagnosis of movement disorders and to monitor disease progression. Although Parkinson’s disease (PD) patients show some oculomotor abnormalities, it is not clear whether previously-described eye movement impairments are task specific. We assessed eye movements in PD under different visual (stationary vs. moving targets) and movement (reactive vs. deliberate) conditions. We demonstrate that PD patients are able to accurately track moving objects but make inaccurate eye movements towards stationary targets. The preservation of eye movements towards dynamic stimuli might enable patients to accurately act upon the predicted motion path of the moving target. These results can inform the development of tools for the rehabilitation or maintenance of functional performance.

---

Eye movements are increasingly used as a clinical tool to enable earlier diagnosis (Marx et al., 2012; De Vos et al., 2020) and to assess disease progression and treatment effects (Patel et al. 2019) in patients with Parkinson’s disease (PD). Cardinal motor symptoms in PD patients include tremor, bradykinesia, and postural instability, but also impairments of oculomotor function (Armstrong, 2008; 2015). Eye movement deficits are especially prevalent when tasks involve higher-level cognitive processing or deliberation, such as remembering the motion path of a target (memory-based pursuit; Fukushima et al., 2015), anticipating or predicting a future sensory event (predictive pursuit; Helmchen et al., 2012; Fukushima et al., 2017), representing more than one concurrent movement goal (double-step task; Bhutani et al., 2013), or exerting executive control over a movement or task (anti-saccades; Chan et al., 2005; Amador et al., 2006). Moreover, PD patients show executive task-dependent deficits, for example, when selecting a target amongst a stream of temporally competing distractors (Zokaei et al., 2020), a process that requires suppressing distracting information, akin to the anti-saccade task.

Many of the fundamental action-regulating functions required for higher-level tasks are mediated to some degree by the basal ganglia (Jenkinson and Brown, 2011; Noorani and Carpenter, 2014), a brain region profoundly affected by degeneration of dopaminergic neurons in the substantia nigra in PD patients (Albin et al., 1989). Aside from their role in oculomotor control (Hikosaka et al., 2000), the basal ganglia might act as a gateway to sensory and memory function (McNab and Klingberg, 2008), as a performance mediator (Thura and Cisek, 2017), and as a key structure involved in sensory evidence accumulation (Perugini et al., 2018) and cancelation of impending actions (Noorani and Carpenter, 2014). Dopaminergic cortical-basal ganglia circuits are implicated in sensory and cognitive deficits in PD patients, especially in situations that require decision making (Perugini et al, 2018).

Despite systematic movement deficits, there appears to be some preservation of motor function in PD patients. For example, “Kinesia Paradoxa” refers to the clinical phenomenon that PD patients perform selected sensory-driven motor tasks with near-normal ability, despite general motor slowing (Glickstein and Stein, 1991; Duysens et al., 2021). In the oculomotor domain, preserved functions include the latency of visually-guided saccades (Briand et al., 1999; Chan et al., 2005) and the initiation of visually-driven smooth pursuit (Fukushima et al., 2015)—functions that are driven by external, visual stimulation (as opposed to self-generated). During reaching, PD patients are able to reach for a moving ball as quickly as controls, but they are impaired when asked to make a self-generated reach for a stationary ball (Majsak et al., 1998). Preserved functions are also found when a movement trajectory has to be corrected online to account for a displacement of the movement target—a task that requires a sense of urgency (Desmurget et al., 2004). Congruently, PD patients performed corrective saccades at a comparable level to healthy controls in a saccade double-step task (Merritt et al., 2017), although they also exhibited a larger number of averaging saccades (Bhutani et al., 2013).

To investigate the accuracy, variability, and preservation of oculomotor functions across different stimuli and task demands, we tested 16 PD patients and 18 healthy, age-matched controls on a battery of movement tasks—pro-saccades, anti-saccades, visually-guided pursuit, and a rapid go/no-go manual interception task. In these tasks, participants viewed either stationary or moving stimuli that elicited reactive or deliberate eye movements (**Figure 1**). The different combinations of stimulus property (stationary vs. moving) and eye movement response (reactive vs. deliberate) allows us to investigate similarities and differences in saccade and pursuit deficiencies as a function of stimulus and task. PD patients showed systematic impairments in tasks that involved stationary targets, indicating impaired saccade inhibition. By contrast, eye and hand movements to moving targets were generally preserved in PD patients as compared to controls.

**Figure 1.**
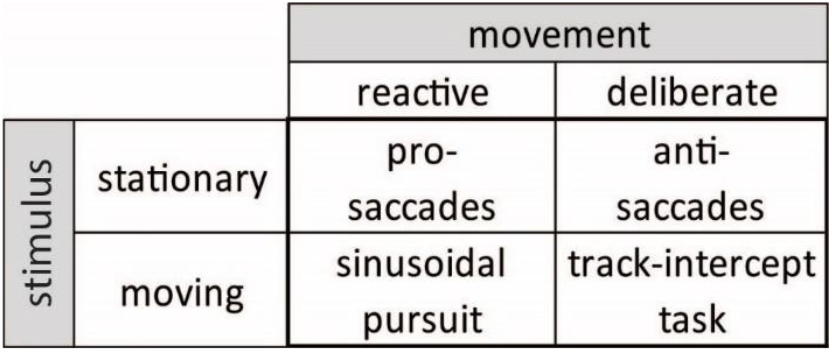
Stimulus characteristics and movement requirements in a battery of oculomotor tasks.

## METHODS

### Participants

Participants were 16 patients with mild to moderate Parkinson’s disease (Hoehn and Yahr 1-2; Goetz et al., 2004) and 18 healthy, age- and sex-matched controls (see **Table 1**). Inclusion criteria for all participants were visual acuity of 20/50 or better, no history of psychiatric or other neurologic disease, including no concussion within the past two years, no history of ocular motility abnormality, and normal cognitive function (Montreal Cognitive Assessment, MoCA, score of 25 or higher). To ensure near-normal visual acuity, all participants were tested using the Early Treatment of Diabetic Retinopathy Study (ETDRS) chart at a 4-m distance (Original Series Chart “R”; Precision Vision, La Salle, IL, USA). Participants with corrective lenses were asked to wear their glasses or contact lenses during testing. All participants confirmed that they were able to clearly see the visual targets. Patients were recruited through the UBC Pacific Parkinson’s Research Centre and affiliated clinical offices and were diagnosed by a neurologist. Controls were recruited from the community. Patients were tested twice, on two different days, once whilst on medication (Levodopa or equivalent; **Table 1**), within two hours of last dose intake, once off medication, after overnight withdrawal of dopaminergic withdrawal; controls were tested once. Testing order for patients (on vs. off medication) was randomized. All experimental procedures were aligned with the Declaration of Helsinki and approved by the University of British Columbia Clinical Research Ethics board; participants gave written informed consent.

**Table 1.** Characteristics of study participants

| Subject Code | Age | Handed-ness | Sex | ETDRS* | MoCA† | Disease Duration (years) | Hoehn-Yahr Stage (0-5)§ | UPDRS Score (0-132)‡ | Dominant Arm Rigidity (0-4) | Test Order | Combination Levodopa\|\| (mg) |
| --- | --- | --- | --- | --- | --- | --- | --- | --- | --- | --- | --- |
| P23 | 67 | RH | M | 20/40-2 | 27 | 3 | 2 | 44 | 2 | N/A | 0 |
| P24 | 78 | RH | F | 20/25-1 | 27 | 6 | 2 | 44 | 1 | ON/OFF | 750 |
| P26 | 84 | RH | M | 20/25-1 | 24 | 14 | 2 | 49 | 3 | ON/OFF | 2250 |
| P29 | 71 | RH | M | 20/20 | 27 | 8 | 2 | 48 | 3 | ON/OFF | 1625 |
| P30 | 61 | RH | F | 20/16-2 | 30 | 9.5 | 2 | 35 | 1 | ON/OFF | 812.5 |
| P31 | 67 | RH | M | 20/16-2 | 27 | 0.5 | 2 | 34 | 3 | ON/OFF | 687.5 |
| P32 | 61 | RH | M | 20/16-1 | 28 | 8 | 2 | 40 | 2 | ON/OFF | 2000 |
| P34 | 65 | RH | M | 20/25-1 | 27 | 4 | 2 | 14 | 1 | ON | 1000 |
| P35 | 78 | RH | F | 20/50-2 | 27 | 16 | 2 | 39 | 2 | ON | 1625 |
| P36 | 67 | RH | M | 20/20-1 | 26 | 10 | 2 | 15 | 0 | ON/OFF | 1000 |
| P37 | 65 | RH | M | 20/25-1 | 28 | 20 | 2 | 29 | 2 | ON/OFF | 750 |
| P38 | 58 | RH | F | 20/20 | 27 | 25 | 3 | 54 | 2 | ON | 1000 |
| P43 | 72 | RH | M | 20/25-2 | 28 | 5 | 2 | 18 | 2 | ON/OFF | 1187.5 |
| P44 | 58 | RH | M | 20/20-1 | 30 | 4 | 2 | 36 | 2 | ON/OFF | 937.5 |
| P45 | 41 | RH | F | 20/12.5-1 | 30 | 3 | 2 | 21 | 2 | ON/OFF | 800 |
| P49 | 70 | RH | M | 20/20-1 | 26 | 13 | 2 | 8 | 0 | ON | 1875 |
| Mean ± SD | 66.4±9.9 |  |  | 20/22-1±0.2 | 27.4±1.6 | 9.71±7.0 | 2.1±0.3 | 33±13.9 | 1.75±0.9 |  | 1143.8±584.0 |
| C25 | 74 | RH | M | 20/25-1 | 26 |  |  |  |  |  |  |
| C27 | 81 | RH | F | 20/16-2 | 28 |  |  |  |  |  |  |
| C28 | 60 | RH | M | 20/32-1 | 25 |  |  |  |  |  |  |
| C39 | 68 | LH | F | 20/20-1 | 28 |  |  |  |  |  |  |
| C40 | 64 | RH | F | 20/20-1 | 30 |  |  |  |  |  |  |
| C41 | 61 | LH | M | 20/25 | 27 |  |  |  |  |  |  |
| C42 | 69 | RH | M | 20/16-1 | 29 |  |  |  |  |  |  |
| C46 | 62 | RH | M | 20/16-2 | 29 |  |  |  |  |  |  |
| C47 | 61 | RH | M | missing | 29 |  |  |  |  |  |  |
| C48 | 74 | LH | M | 20/12.5-2 | 28 |  |  |  |  |  |  |
| C50 | 69 | RH | F | 20/20-1 | 26 |  |  |  |  |  |  |
| C51 | 78 | RH | M | 20/20-2 | 26 |  |  |  |  |  |  |
| C52 | 71 | RH | M | 20/25-1 | 28 |  |  |  |  |  |  |
| C53 | 69 | RH | M | 20/16-1 | 29 |  |  |  |  |  |  |
| C54 | 79 | RH | M | 20/20 | 30 |  |  |  |  |  |  |
| C55 | 88 | RH | M | 20/25 | 28 |  |  |  |  |  |  |
| C56 | 65 | RH | M | 20/50-1 | 30 |  |  |  |  |  |  |
| C57 | 43 | RH | F | 20/20 | 30 |  |  |  |  |  |  |
| Mean ± SD | 68.7±10.0 |  |  | 20/22±0.2 | 28.1±1.6 |  |  |  |  |  |  |

\* Early Treatment of Diabetic Retinopathy (ETDRS) visual acuity chart “R” (Precision Vision).
† Montreal Cognitive Assessment, a test that rates cognitive ability on a scale from 0 to 30 (Nasreddine et al. 2005)
§ Hoehn and Yahr (1967) staging scale for symptom severity, ranging from 1 (unilateral involvement only) to 5 (confinement to bed or wheelchair).
‡ Unified Parkinson’s Disease Rating Scale (Movement Disorder Society Task Force 2003). Motor Score only.
|| Most patients were on combination drugs containing Levodopa and Carbidopa (e.g., Sinemet, Levocarb). Table states total daily dose in milligram (mg) across equivalent combination drugs.

### Visual Display and Apparatus

Stimuli were back-projected onto a translucent screen with a PROPixx video projector (VPixx Technologies, Saint-Bruno, QC, Canada; refresh rate 60 Hz, resolution 1,280 (horizontal) × 1,024 (vertical) pixels. The displayed window was 40.7 (horizontal) × 33.3 (vertical) cm or 67 degrees of visual angle [°] × 60° in size. Stimulus display and data collection were controlled by a PC (NVIDIA GeForce GT 430 graphics card) and the experiment was programmed in MATLAB 7.1 using Psychtoolbox 3.0.8 (Brainard 1997; Kleiner et al. 2007; Pelli 1997). Participants were seated in a dimly-lit room at 46 cm distance from the screen with their head supported by a combined chin and forehead rest.

### Saccade and pursuit tasks

Participants first performed a pro- and anti-saccade task (Munoz and Everling, 2004), designed to test saccade control at different levels of deliberation (**Fig. 1**). Pro and anti-saccade targets were presented on a black background (0.06 cd/m^2^). The pro-saccade task (**Fig. 2A**) started with a green fixation square (0.8° side length; 69.7 cd/m^2^) shown at the screen centre; eye tracker drift correction was performed during initial fixation. At the same time as the fixation square, two white target squares (each 0.8°; 96.5 cd/m^2^) were presented in the periphery, at 12° to the left and right of fixation. After a random fixation period (0.8-1.2 s) an open square (1.2° side length) appeared around one of the white target squares, indicating the side to which participants should move their eyes. The offset of the green fixation square served as a cue to initiate a saccade toward the target. The anti-saccade task (**Fig. 3A**) followed the same timeline, except that here, the fixation square was red (0.8° side length; 21.6 cd/m^2^), and the open square marked the distractor, i.e., participants had to look away from it and toward the uncued target. Each participant completed 40 trials of each task.

**Figure 2.**
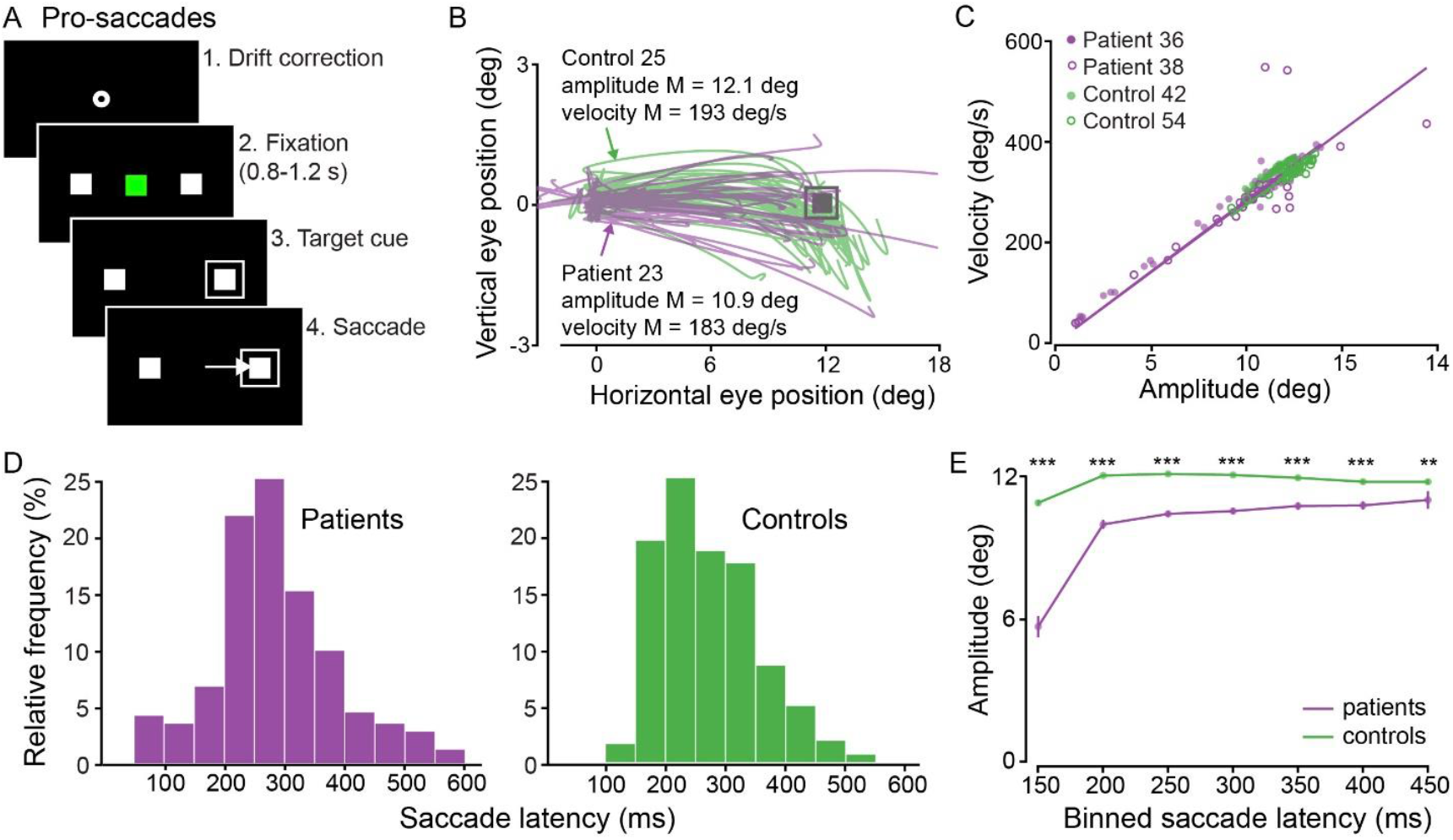
Sequence of events and eye movements in the pro-saccade task. (A) Each trial started with a drift correction followed by a fixation period. Participants had to saccade to the cued target square. (B) 2D eye position in pro-saccade task for a representative PD patient (purple) and control participant (green). For illustration purposes, eye and target position data were flipped to always depict the saccade target on the right. (C) Main sequence (saccade velocity vs. amplitude) for two representative patients (purple circles) and two control participants (green circles). Each circle represents one trial. (D) Saccade latency distributions (relative frequency of binned saccade latencies) for patients and controls. (E) Mean saccade amplitude as a function of saccade latency. Each dot represents the mean saccade amplitude in a 50 ms time bin across all patients (purple) and controls (green). Vertical lines indicate standard error. Asterisks denote significance level of ranked sum test: ** *p* < .01 and *** *p* < .001.

**Figure 3.**
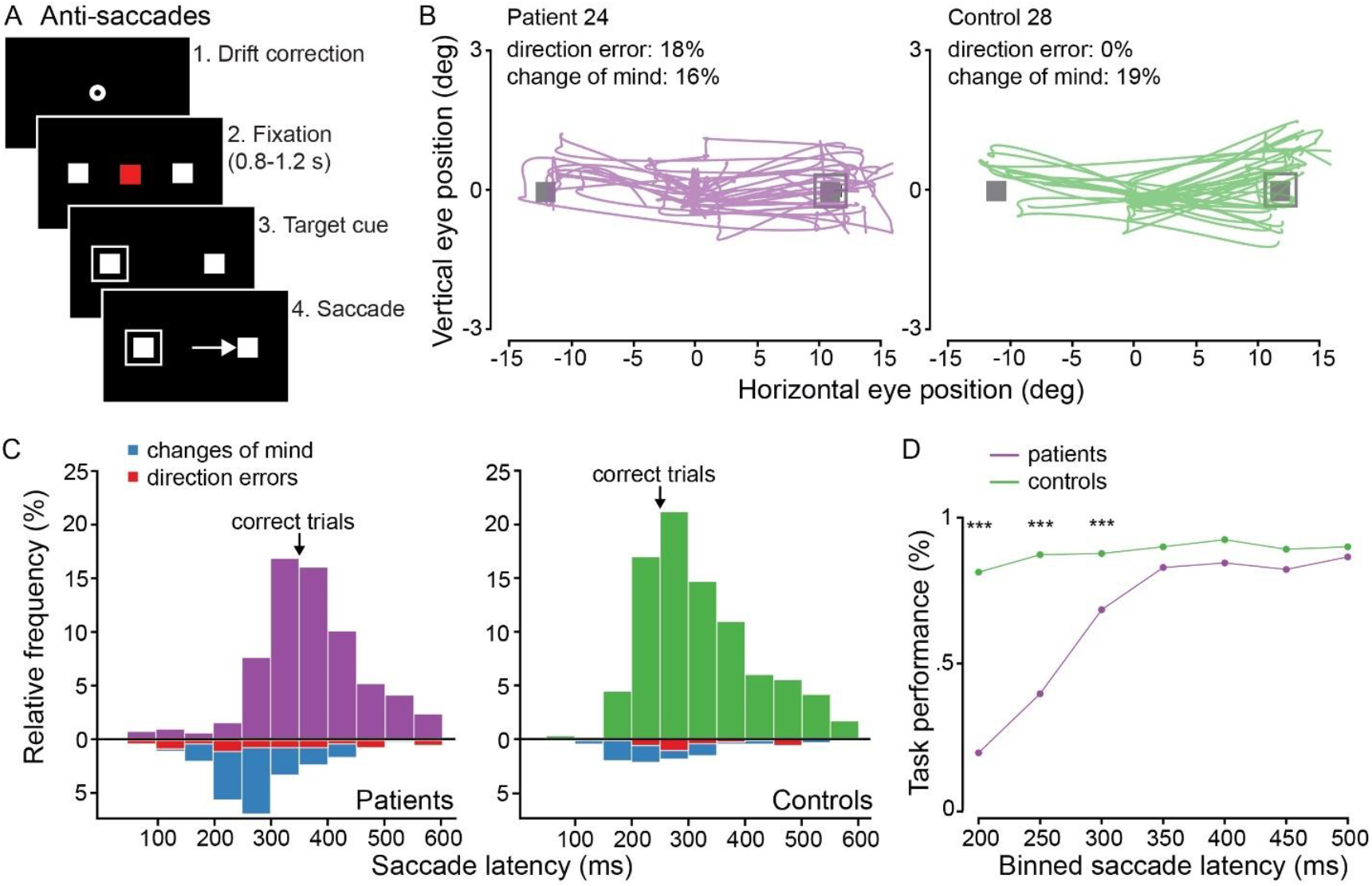
Sequence of events and eye movements in the anti-saccade task. (A) Each trial started with a drift correction followed by a fixation period. Participants had to saccade to the uncued target square. (B) 2D eye position in pro-saccade task for a representative PD patient (purple) and control participant (green). For illustration purposes, eye and target position data were flipped to always depict the saccade target on the right. (C) Saccade latency distributions (relative frequency of binned saccade latencies) for patients and controls. Blue bins indicate changes of mind and red bins indicate direction errors. (D) Task performance (percentage of saccades towards uncued location without any corrections) as a function of saccade latency. Asterisks denote significance level of ranked sum test: *** *p* < .001.

Participants next performed a baseline smooth pursuit tracking task. This task was designed to characterise basic tracking function akin to testing pursuit at the bedside by regularly moving a small object to-and-fro at different speeds before the patient’s eyes (Leigh and Zee, 2015). Each trial started with a drift correction (fixation on a central bull’s eye stimulus 2° in diameter). The smooth pursuit target was a small (2° in diameter) black disk presented on a grey background with a luminance of 97.6 candela per meter squared (cd/m^2^). The target moved sinusoidally for five repetitions at 16°/s or 32°/s, first along the horizontal and then along the vertical meridian (**Fig. 5A**). Reflection points were positioned at ± 16° to the left/right or top/down and each speed was presented once per motion direction resulting in 4 trials per participant.

### Track-intercept task

In the second part of testing, participants performed a timed go/no-go task, in which they had to track and manually intercept a moving target that followed a linear-diagonal path and either hit or missed a dedicated strike box (**Fig. 6A**). The moving target was a black Gaussian dot (SD = 0.4°; d = 2°; 5.4 cd/m^2^) presented on a gray background (35.9 cd/m^2^). The strike box (31.5 cd/m^2^) was 6° × 10° in size and offset by 12° from the center to the side of interception. Importantly, the target was only shown for 300 or 500 ms and then disappeared. Participants had to predict whether the target would pass or miss the strike box by following the target’s assumed trajectory even after it had disappeared. They were asked to intercept the target while it was in the strike box in pass trajectories, and withhold a hand movement in miss trajectories. Each interception started from a table-fixed position and was made with the index finger of the dominant hand. Stimulus velocity followed natural forces (gravity, drag force, Magnus force; Fooken and Spering, 2019). The target launched at an angle of 5°-12°, depending on the type of trajectory, and moved at a speed of either 13 or 17°/s; conditions were presented in randomized order. Each trial ended when participants either intercepted the target or when the target reached the edge of the screen (2-2.6 s). At the end of each trial participants received performance feedback; target end position was shown, and correct or incorrect decisions were indicated. Each participant performed a familiarization session (8 trials; full trajectory visible) followed by 120 experimental trials in which the target viewing time was limited.

### Eye and hand movement recordings and preprocessing

Eye position of the right eye was recorded with a video-based eye tracker (Eyelink 1000 tower mount; SR Research Ltd., Ottawa, ON, Canada) at a sampling rate of 1000 Hz. Eye movements were analyzed off-line using custom-made routines in MATLAB (R2015a). Eye velocity profiles were filtered using a low-pass, second-order Butterworth filter with cut-off frequencies of 15 Hz (position) and 30 Hz (velocity). Saccades were detected based on a combined velocity and acceleration criterion: five consecutive frames had to exceed a fixed velocity criterion of 30°/s; saccade on- and offsets were then determined as acceleration minima and maxima, respectively. Saccades were excluded from smooth pursuit analysis. Pursuit onset was detected in individual traces using a piecewise linear function that was fit to the filtered position trace.

Finger position was recorded with a magnetic tracker (3D Guidance trakSTAR, Ascension Technology Corp., Shelburne, VT, USA) at a sampling rate of 60 Hz; a lightweight sensor was attached to the participant’s dominant hand’s index fingertip with a small Velcro strap. Finger latency was defined as the first sample in which finger velocity exceeded 5% of the finger’s peak velocity. The 2D finger interception position was recorded in x- and y-screen-centered coordinates.

### Eye and hand movement performance measures

For all eye and hand movement measures reported in the manuscript we calculated an average value per participant by finding the median value across trials. We also assessed within-participant variability by calculating the standard deviation of a given measure across trials. We aimed to test patients on two separate visits when they were either on or off their medication (counterbalanced order). Four patients were unable to come in for testing while off medication and one patient did not take any medication (P23, **Table1**). For the remaining 11 patients we found no effect of medication on eye movement timing and accuracy (e.g., saccade amplitude, *t*(23.5)=3.90, p<.001, in the pro-saccade task or on sensorimotor decision accuracy, *t*(24.9)=1.26, p=.22). Because patients generally had noisier data than controls we had a higher rate of trial exclusions in patients (see below). Therefore, we decided to pool data from both test days for all patients who came in twice (unless reported otherwise). To ensure that unequal trial numbers across participants did not affect our main results we repeated each analysis using only data from the first visit. These results did not statistically differ from the results reported here.

Saccade performance in the pro- and anti-saccade task was quantified by calculating saccade latency, velocity, duration, and amplitude. Saccade latency was defined as the difference between target cue and first saccade onset. Saccades with a latency of <150 ms were defined as express saccades (Fischer, 1987). We then determined the velocity, duration, and 2D amplitude of this initial saccade. For the anti-saccade task, we also calculated the number of direction errors (i.e., saccades directed to the cued rather than uncued target and not later corrected) and the number of changes of mind (i.e., saccades initially directed to the cued target, but then corrected to the uncued target).

Smooth pursuit accuracy was quantified by calculating pursuit latency, gain, position error, and saccade rate. Pursuit latency was defined as the time difference between stimulus onset and pursuit onset. If no pursuit was initiated and participants fixated until initiating a saccade, pursuit onset was defined as the offset of that first saccade. The rate of catch-up saccades was defined as the average number of saccades per second across the entire trial. Pursuit gain, eye position error and catch-up saccade rate were analysed during steady-state pursuit, omitting the response within 140 ms of either side of the target deflection points. Gain was defined as the mean relative difference between eye and target velocity; eye position error was defined as the 2D distance between eye and target position. Pursuit gain and eye position error were calculated during smooth tracking (excluding saccades and blinks).

For the track-intercept task we calculated pursuit latency, initial eye velocity, horizontal position error and saccade rate while the target was visible (300 or 500 ms), and the latency of the first catch-up saccade. For the finger, we analyzed finger latency, peak velocity, interception timing error, and positional interception error. Finger latency was defined as the difference between target onset and finger movement onset. Interception timing error was calculated by dividing the distance between the target and the point of interception by the average target velocity. Positional interception error was calculated as the 2D error between target position and hand position at time of interception. To calculate hand movement speed adjustment within an experimental session we used the first session for patients that were tested on and off medication.

All trials were manually inspected and trials, in which participants blinked during target presentation were excluded from analysing the given task. Based on inspection, we excluded one participant for the pro- and anti-saccade task because no valid eye movement data were collected. We also excluded four control subjects from the manual interception task that had more than 25% trials of eye movement signal loss. Following the same cut-off (more than 25% of invalid trials), we also excluded data from one patient on ON-medication day and data from two patients on OFF-medication day. Usable data from the respective other testing days were included in the analysis.

For the remaining participants, we excluded 132 trials (1%) in the pro-saccade task, 159 trials (1%) in the anti-saccade task, and 575 (12%) in the manual interception task.

### Statistical analyses

Differences between PD patients and controls were evaluated using Welch’s two-sample unpaired *t*-tests. We used Welch’s t-tests to adjust for the variance *S* of each group of size *N*. Degrees of freedom using Welch t-tests are estimated as follow

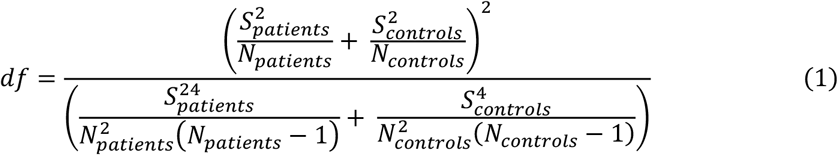

Pooled group differences for saccade latency dependent intervals were compared using a Mann-Whitney test. We assessed the probability of group values being not equal (*p* value) and the z-Score (*z* value). A *z* value close to 0 indicates that group medians are equal. To compare oculomotor performance across tasks we calculated a linear regression and correlation coefficient. All statistical analyses were performed using R (version 4.01, R Core Team, 2017).

## RESULTS

Early-stage PD patients with mild to moderate symptoms and age-matched healthy controls performed a variety of movement tasks that required sensorimotor decisions at different levels of task complexity. The tasks ranged from visually guided pro- and anti-saccades, baseline smooth pursuit tracking, to rapid go/no-go manual interceptions.

### Eye movements to stationary targets are impaired in PD patients

In the first part of the experiments, participants were instructed to quickly move their eyes either to a stationary target that was cued (pro-saccades) or to a stationary target that was located opposite to a cued distractor (anti-saccades). In both tasks we found systematic differences in eye movement speed, accuracy, and variability between patients (pooled across ON and OFF medication) and controls. In the pro-saccade task (**Fig. 2A**), patients tended to undershoot the saccade target on average (i.e., saccades were hypometric), whereas controls landed on the target on average (**Fig. 2B**). Moreover, patients’ saccades were slower (lower peak velocity) as compared to controls (**Table 2**). To investigate whether the velocity reduction in patients’ saccades was linked to their saccade hypometria, we considered the relationship between saccade velocity and amplitude (main sequence; **Fig. 2C**). We found that patients and controls showed a positive linear relationship between saccade velocity and amplitude with comparable slopes (*M*_*patients*_ = 22.8 ± 5.0 1/s; *M*_*controls*_ = 25.0 ± 4.7 1/s; *t*(32) = 1.33, *p* = .19). These findings indicate that slower saccades in patients could be linked to the fact that their saccades are also of smaller size. Whereas the general relationship between saccade velocity and amplitude was comparable between patients and controls, we found that patients’ saccades were more variable across trials (see examples in **Fig. 2C**). This within-participant eye movement variability was reflected in significantly higher standard deviations of saccade amplitude, velocity, and latency in patients as compared to controls (**Table 3**).

**Table 2.**
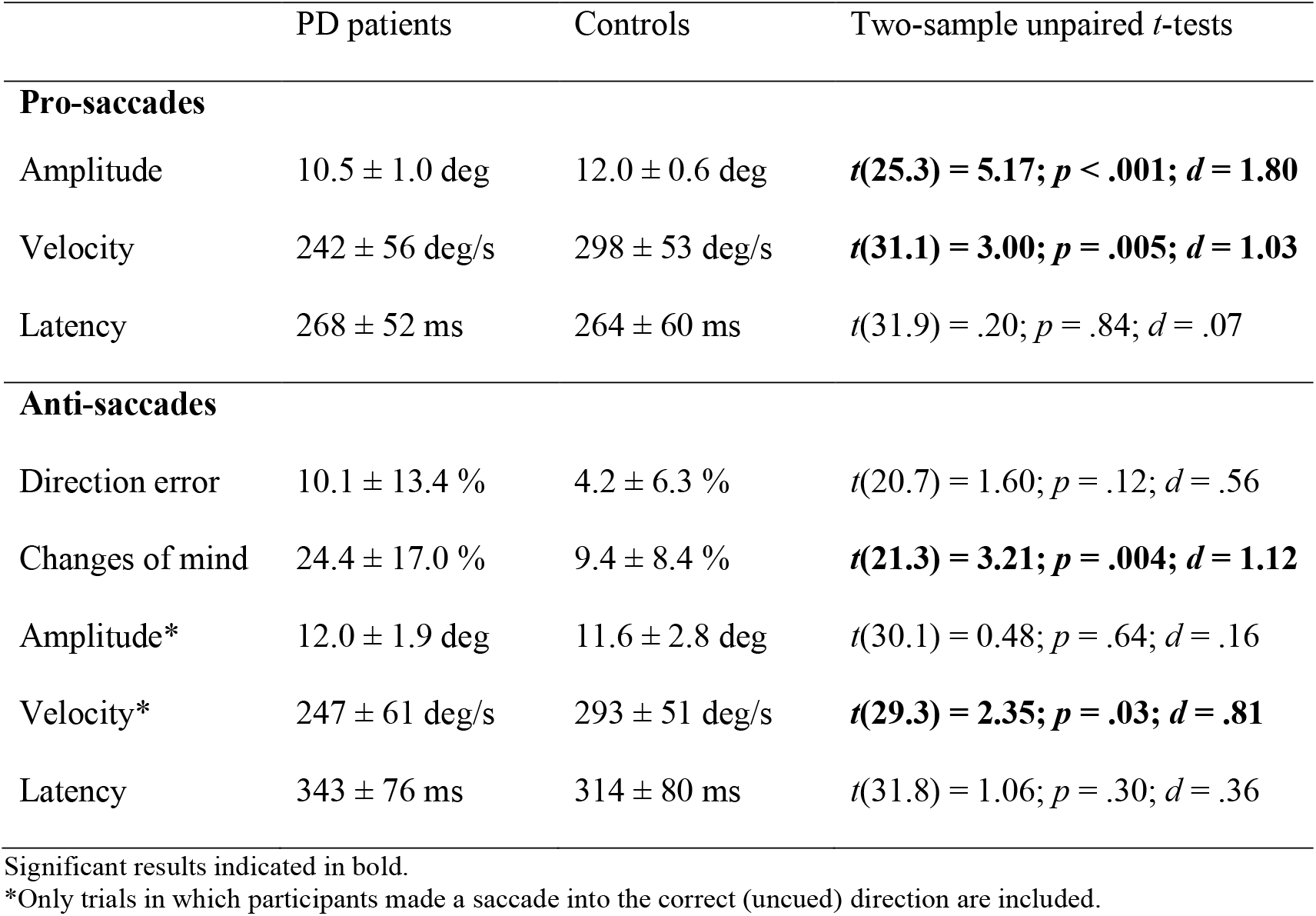
Saccadic eye-movement accuracy.

**Table 3.** Saccadic eye-movement variability.

|  | PD patients | Controls | Two-sample unpaired *t*-tests |
| --- | --- | --- | --- |
| **Pro-saccades** |  |  |  |
| Amplitude | 3.3 ± 1.5 deg | 0.8 ± 0.4 deg | ***t*(16.9) = 6.45; *p* < .001; *d* = 2.27** |
| Velocity | 72 ± 34 deg/s | 29 ± 21 deg/s | ***t*(24.4) = 4.40; *p* < .001; *d* = 1.53** |
| Latency | 106 ± 35 ms | 55 ± 21 ms | ***t*(24.3) = 5.14; *p* < .001; *d* = 1.79** |
| **Anti-saccades** |  |  |  |
| Amplitude* | 3.1 ± 2.0 deg | 1.5 ± 1.2 deg | ***t*(23.8) = 2.83; *p* = .009; *d* = 0.99** |
| Velocity* | 66 ± 44 deg/s | 30 ± 19 deg/s | ***t*(19.8) = 3.01; *p* = .007; *d* = 1.06** |
| Latency | 115 ± 29 ms | 73 ± 26 ms | ***t*(30.5) = 4.45; *p* < .001; *d* = 1.53** |

Significant results indicated in bold.
\*Only trials in which participants made a saccade into the correct (uncued) direction are included.

In the pro-saccade task, saccade latencies ranged from 50-600 ms (**Fig. 2E**). Notably, patients made more express saccades with latencies shorter than 150 ms compared to controls (patients: 7.8%; controls: 1.7%). To investigate whether increased latency variability in patients could be linked to saccade accuracy, we analyzed saccade amplitude as a function of saccade latency at a group level. Overall, saccades were hypometric (inaccurate) in patients compared to controls for all latency intervals (*p<*.001 and z>3.62 for all latencies shorter than 450 ms and *p*=.004 and *z*=2.91 for latencies longer than 450 ms). Interestingly, hypometric saccades in patients were particularly prominent at the shortest saccade latency interval (**Fig. 2E**). These results suggest that patients might have made reflexive saccades toward the cued target before motor planning was complete.

In the anti-saccade task, participants had to inhibit a saccade response to a cued distractor location and instead make a deliberate saccade to the opposite side (**Fig. 3A**). We assessed task performance by describing two types of errors: direction errors are defined as saccades that landed on the cued target location and were not subsequently corrected. Changes of mind are defined as saccades that were initially directed to the cued target location but then corrected to the opposite side. In patients and controls, the frequency of direction errors was lower than the frequency of changes of mind, indicating that most saccades that were initially directed at the cued distractor were subsequently corrected (**Table 2**). Overall, patients made about twice as many errors as controls, and were significantly more likely to change their mind as compared to controls (**Fig. 3B**; **Table 2**).

Similar to the pro-saccade task, we observed that patients had more variable eye movement amplitudes, velocities, and latencies across trials (within-participant variability) compared to controls (**Table 3**). We compared saccade kinematics for trials in which participants correctly performed the task (excluding trials with direction errors and changes of minds). As in the pro-saccade task, patients made slower saccades than controls (**Table 2**), but anti-saccades were overall of similar amplitude in both groups of participants (**Fig. 3B**). These findings indicate that hypometria might overall be less prevalent in a task that required more deliberation and triggered longer saccade latencies as compared to a visually-cued saccade task.

We next evaluated task performance (correct trials, direction errors and changes of mind) as a function of saccade latency. Even though patients initiated saccades at around the same time as controls (**Table 2**), their task performance depended on saccade latencies. Shorter saccade latencies were associated with more errors (**Fig. 3C-D**), in fact, patients only made more errors than controls for saccades with latencies shorter than 300 ms (*p* < .001 and *z* > 5.15). These findings mirror the observation that short-latency pro-saccades in patients tend to be hypometric and indicate that patients’ saccade task performance in generally is most impaired for short-latency saccades.

To directly link performance in the pro- and anti-saccade task we chose two measures that were indicative of task performance and were related to successful saccade inhibition. For the pro-saccade task, we calculated the percentage of express saccades participants made towards the cued target. For the anti-saccade task, we used the frequency of task errors (direction errors and changes of mind). We then related these performance measures across tasks. In the patient group, we found a positive correlation (*r* = .85) between express saccades in the pro-saccade task and task error rate in the anti-saccade task (**Fig. 4A**). No such relationship was found in the control group. Only one control participant (C57; a highly-trained vision scientist who is one of the authors) initiated saccades with latencies shorter than 150 ms, but her task error rate was low. Comparing saccade latency distributions between C57 and a PD patient that had the same rate of express saccades (P35) illustrates a key difference. Whereas C57 has a narrow distribution of saccades centered around a latency of approximately 175 ms, P35 has an initial distribution of express saccades that peaks around 75 ms and then another wide-spread distribution of longer-latency saccades (left panel in **Fig. 4B**). The observation that the rate of express saccades during the pro-saccade task was linked to the rate of errors during the anti-saccade task in PD patients suggests that eye movements to stationary targets are controlled similarly irrespective of the level of movement deliberation.

**Figure 4.**
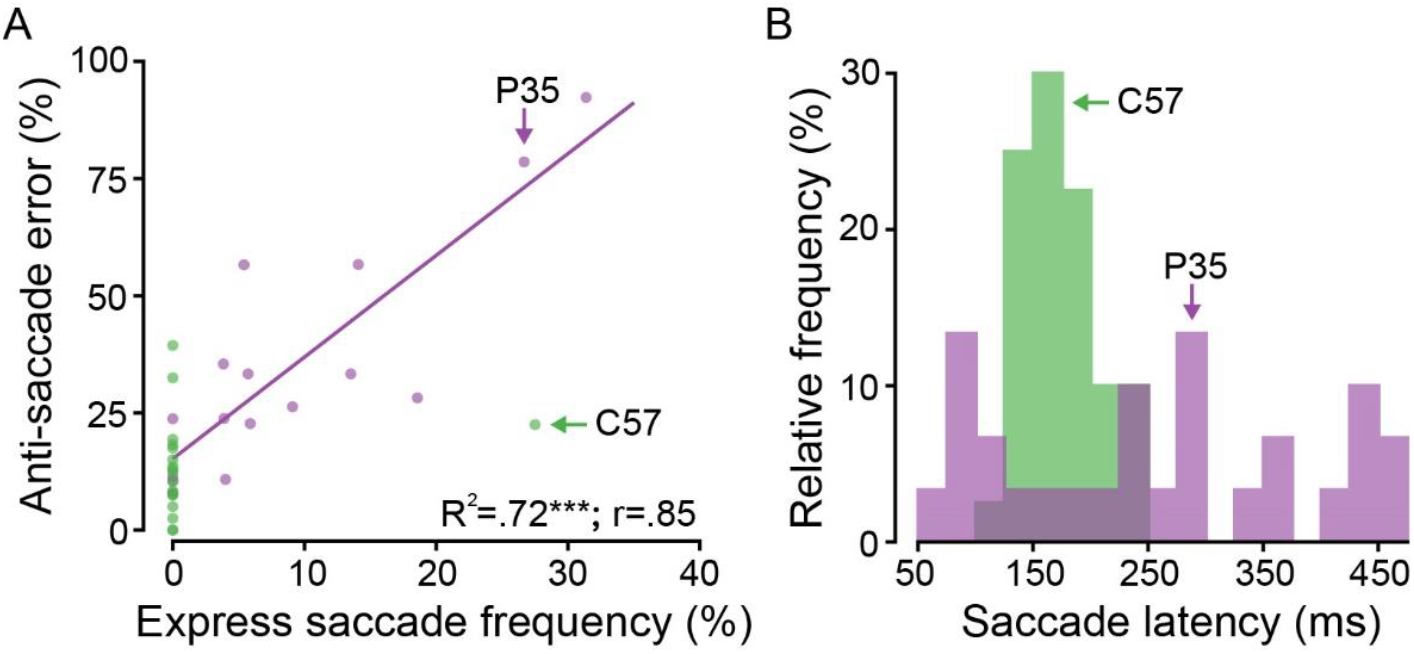
Comparison of pro- and anti-saccade task performance. (A) Relationship between the frequency of express saccades during the pro-saccade task and the error rate (saccade towards the cued target) in the anti-saccade task. Each circle represents a patient (purple) and control participant (green). Asterisk denotes significant regression results in patient group: \*\*\**p <* 0.001. (B) Saccade distributions of a control participant (C57; green) and patient (P35; purple) who had a similar rate of express saccades.

### Eye and hand movements to moving targets are preserved in PD patients

Participants performed two tasks that involved moving targets. In the baseline pursuit task, participants were asked to follow a moving target with their eyes; in the go/no-go track-intercept task participants had to follow and manually intercept a moving target that disappeared after brief initial presentation. In the baseline pursuit task (**Fig. 5A**), we found that patients were able to track the moving target with similar speed and accuracy as controls (**Fig. 5B**). Even though patients made more catch-up saccades on average to keep their eyes aligned with the moving target, patients’ saccades during pursuit were as accurate as controls’ (comparable position error) indicating that pursuit performance was overall preserved (**Table 4**).

**Figure 5.**
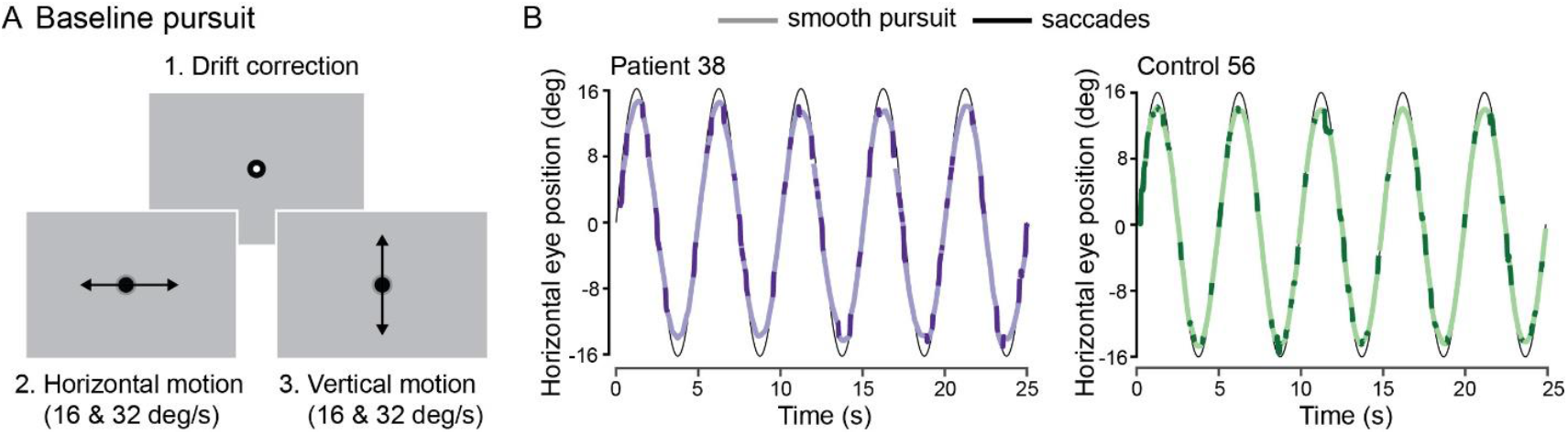
Sequence of events and eye movements in baseline pursuit task. (A) Each trial started with a drift correction followed by five cycles of sinusoidal target motion in either horizontal or vertical direction. (B) 2D eye position for a horizontally moving target at a speed of 16 deg/s for a representative PD patient (purple) and control participant (green). Saturated segments denote saccades, lighter segments represent smooth pursuit.

**Table 4.** Eye movement accuracy during baseline pursuit and track-intercept task.

|  | PD patients | Controls | Two-sample unpaired *t*-tests |
| --- | --- | --- | --- |
| **Smooth pursuit** |  |  |  |
| Eye velocity gain | 1.08 ± .23 | 1.01 ± .21 | $t(30.5) = .87$; $p = .39$; $d = .30$ |
| Position error | 2.2 ± 1.0 deg | 1.9 ± .7 deg | $t(25.4) = .79$; $p = .44$; $d = .27$ |
| Saccade rate | 4.6 ± 1.1 sac/s | 4.0 ± .7 sac/s | **$t(24.4) = 2.05$; $p = .05$; $d = .71$** |
| **Track-intercept** |  |  |  |
| Pursuit latency | 88 ± 48 ms | 49 ± 51 ms | **$t(26.8) = 2.14$; $p = .04$; $d = .79$** |
| Initial eye velocity | 5.8 ± 1.6 deg/s | 6.1 ± 1.6 deg/s | $t(27.2) = .46$; $p = .65$; $d = .17$ |
| Position error | 1.3 ± .3 deg | 1.2 ± .3 deg | $t(26.1) = 1.47$; $p = .15$; $d = .54$ |
| Saccade latency | 275 ± 32 ms | 241 ± 26 ms | **$t(27.9) = 3.18$; $p = .004$; $d = 1.16$** |

During the go/no-go track-intercept task, participants viewed a moving target that disappeared after 300 or 500 ms before passing through or missing an indicated strike zone (**Fig. 6A**). In each trial, participants had to predict whether the no longer visible target would pass (go response required) or miss (no-go required). We first compared how well participants were able to track the moving target with their eyes while it was visible. Similar to baseline pursuit, we found that patients’ tracking was as fast and as accurate as controls’ pursuit, with comparable eye velocity and position errors (**Fig. 6B, Table 4**). However, patients initiated smooth pursuit later and made their first catch-up saccade toward the target later than controls (**Fig. 6C**), indicating that patients showed less anticipation of predictable target motion. Notwithstanding these differences in eye movement timing, patients’ go/no-go decision accuracy—i.e., correctly differentiating whether the target would hit or miss the strike zone—was similar to performance in controls (*M*_*patients*_ = 79.2%, *M*_*controls*_ = 83.7%; *t*(27.7) = 1.12; *p* = .27; *d* = .41). Because we found performance differences as a function of saccade latency in our saccade tasks, we next analyzed go/no-go decision accuracy on a group level as a function of the first saccade latency. We find that patients have less early catch-up saccades compared to controls (**Fig. 6C**). However, congruent with findings in the pro-saccade and anti-saccade tasks, patients were relatively less accurate in their go/no-go decisions compared to controls when initial catch-up saccades were shorter than 150 ms (*p* = .004; *z* = 2.92).

**Figure 6.**
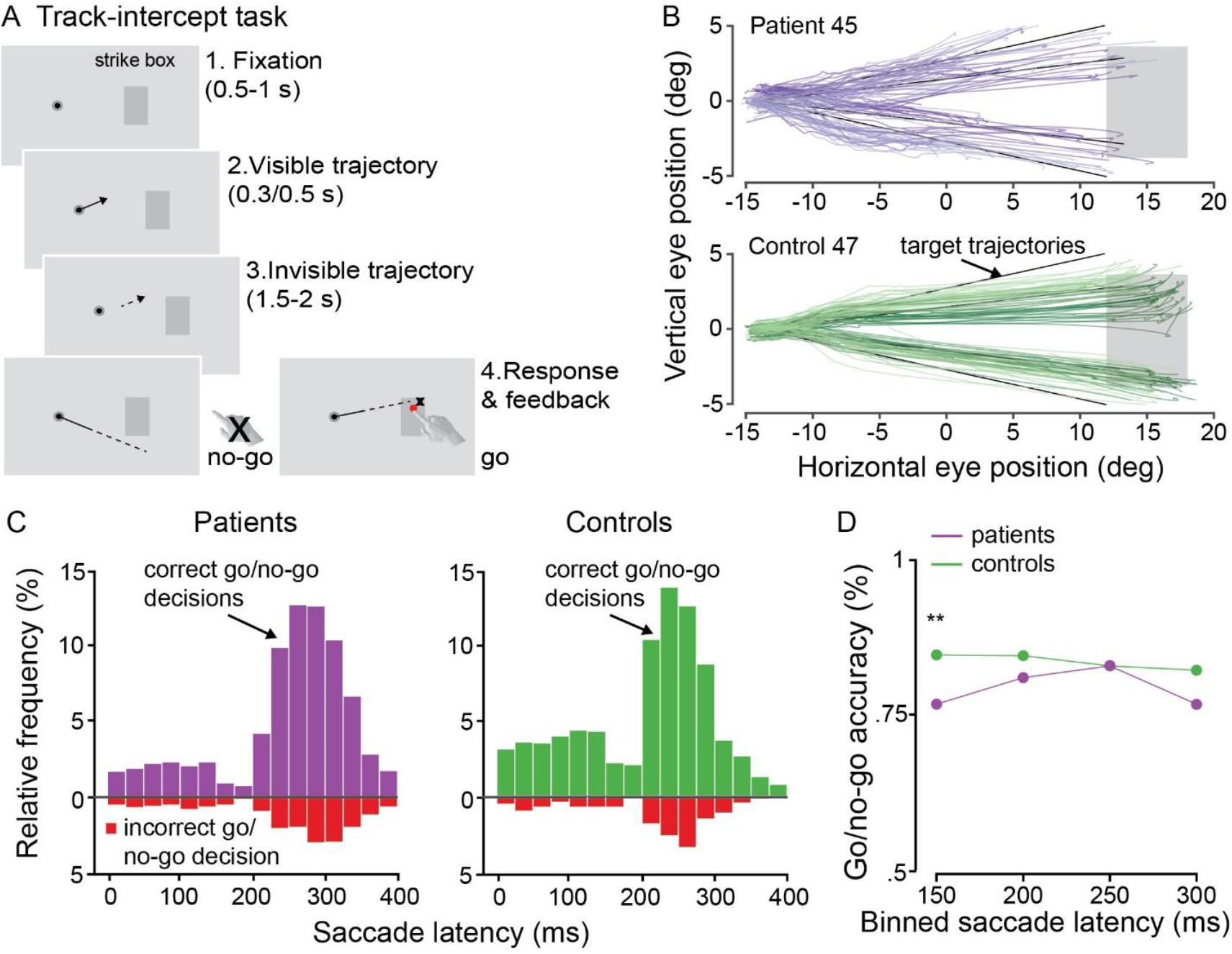
Sequence of events and eye movements in go/no-go track-intercept task. (A) Each trial started with a fixation period. Participants viewed a moving, disappearing target and had to judge whether the target would miss or pass a strike box. (B) 2D eye position in track-intercept task for a representative PD patient (purple) and control participant (green). (C) First catch-up saccade latency distributions (relative frequency of binned saccade latencies) for patients and controls. Red bins indicate trials in which the go/no-go decision was incorrect. (D) Go/no-go decision accuracy as a function of initial catch-up saccade latency for patients (purple) and controls (green). Circles indicate group mean for given saccade interval. Two asterisks denote significance level *p* < .01 of ranked sum test.

### Hand movement deficits are compensated during track-intercept task

The go/no-go track-intercept task required a decision of whether to initiate or withhold a hand movement. Following a go-decision, participants had to move their hand to the strike box and intercept the moving target at the right time. A comparison of hand movement dynamics showed that patients moved their hand slower on average than controls (**Fig. 7A**). However, patients initiated their hand movement ∼150 ms earlier than controls (**Table 5**). These results suggest that PD patients might have compensated for hand movement deficits, such as motor slowing, by starting the interceptive hand movement earlier than controls. Notwithstanding these differences in hand movement latency and velocity between patients and controls, both groups intercepted the target with a comparable timing error—100 ms too early on average (**Fig. 7B**)—and overshot the target location with the same average interception error (**Table 5**). These findings show that interception timing and accuracy are preserved in PD patients despite motor slowing.

**Figure 7.**
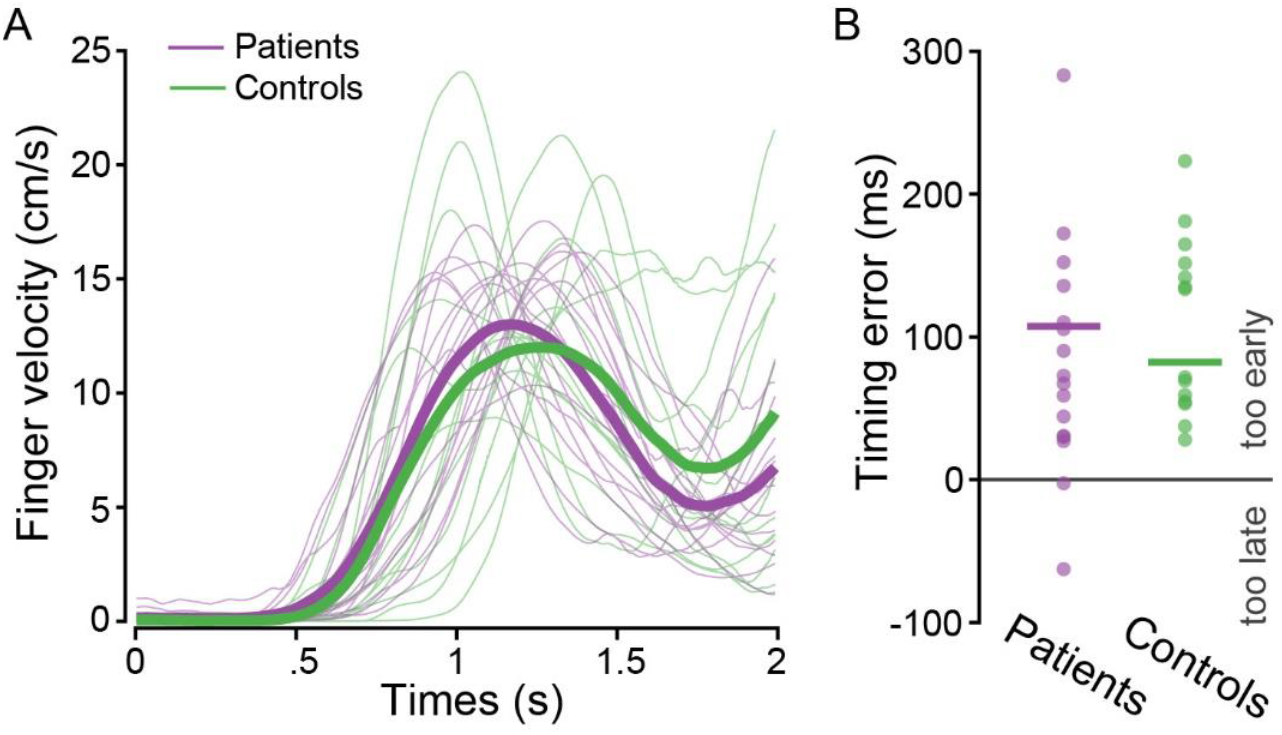
Hand movement dynamics in track-intercept task. (A) Hand movement velocity across time for individual (thin lines) patients (purple) and controls (green). Thick lines represent group average. (B) Interception timing error for patients and controls. Positive timing errors indicate that participants intercepted too early, negative timing error indicate late interceptions.

**Table 5.** Hand movement kinematics during track-intercept task.

|  | PD patients | Controls | Two-sample unpaired *t*-tests |
| --- | --- | --- | --- |
| Latency | 712 ± 155 ms | 868 ± 199 ms | *t*(24.5) = 2.38; *p* = .03; *d* = .88 |
| Peak velocity | 25.6 ± 4.7 cm/s | 32.0 ± 8.1 cm/s | *t*(20.2) = 2.55; *p* = .02; *d* = .95 |
| Interception error | 4.4 ± 1.6 deg | 4.4 ± 1.2 deg | *t*(27.1) = 0.13; *p* = .90; *d* = .05 |

## Discussion

Oculomotor function is known to be systematically impaired in patients with Parkinson’s disease. Here we argue against a general oculomotor decline and show instead that oculomotor deficits are strongly stimulus and task dependent. Our findings provide evidence for differential vulnerability for oculomotor responses to stationary vs. moving stimuli. Different pathologic disease processes might underlie functional decline in response to different types of visual stimulation. In summary, we report the following key findings.

Patients showed systematic impairments when making saccades to stationary targets, regardless of whether the task required reactive pro-saccades or more deliberate anti-saccades. Patients’ pro-saccades were hypometric and anti-saccades went the wrong direction more frequently than for controls. In patients, pro-saccade accuracy and anti-saccade direction errors were more pronounced when saccade latencies were short, suggesting that impairments on both tasks are due to common mechanisms. Overall, patients had difficulties inhibiting reactive saccades to a cued target or distractor, leaving less time to complete accurate motor planning.

Patients did not show impairment when tracking a moving object using a combination of smooth pursuit and saccades. Although patients made more catch-up saccades than controls during baseline pursuit, we did not observe any differences in eye position error or pursuit velocity gain. These results suggest that eye movements to moving stimuli are relatively preserved in PD. Congruently, we found that patients were able to accurately track and predict the trajectory of a moving target that disappeared after a brief viewing time. Go/no-go decision accuracy and timing were overall preserved in patients, except when they initiated a very early catch-up saccade toward the target, thereby limiting time for sensory evidence accumulation. Patients moved their hand slower than controls but were able to compensate by initiating their movements earlier, potentially indicating a learned adjustment to changes in motor function.

### Differential vulnerability to stationary vs. dynamic visual stimulation

In recent years, saccade tasks have become a useful clinical tool to investigate the control and inhibition of eye movements towards visual stimuli in psychiatric and neurological patient populations (Everling and Fischer, 1998; Hutton and Ettinger, 2006; Patel et al., 2019). In PD, saccades toward stationary (visual or remembered) targets are hypometric (Rottach et al., 1996; Gurvich et al., 2007; Helmchen et al., 2012), presumably due to excessive SC inhibition (Terao et al., 2011). In anti-saccade tasks, patients make more incorrect saccades toward the distractor and exhibit a higher saccade latency than controls (Briand et al, 1999; Chan et al., 2005; Amador et al., 2006; for a review, see Waldthaler et al., 2020). Our study adds to these findings by showing that task-specific errors (hypometric pro-saccades, incorrect anti-saccades) occurred predominantly in short-latency saccades. We interpret this finding as evidence of incomplete motor planning: if a saccade is made early, there is less time for accurate direction and endpoint planning (Viviani & Swensson, 1982; Findlay, 1983; Cameron et al., 2012). Both the increase in error rate in the anti-saccade task and the increase in express saccades during the pro-saccade task suggest that PD patients demonstrate decreased inhibitory control (see also Ouerfelli-Ethier et al., 2018 for across-task dependencies). Deficits in inhibitory control might not only be related to impairments in oculomotor pathways but could also be the consequence of adaptive motor control. To counteract slow movement initiation (commonly observed in PD patients) the oculomotor system might reduce baseline response inhibition (Chan et al., 2005). Here we show that PD patients were, in fact, able to initiate an interceptive hand movement towards a moving target earlier than controls. These findings suggest long-term adaptive mechanisms that could be related to an altered baseline response inhibition.

An impairment of movement towards stationary targets is also observed during reaching. Whereas PD patients exhibited bradykinesia when reaching for a stationary object, they moved as fast as controls and with comparable accuracy when reaching for a moving object (Majsak et al., 1998; 2008). These studies highlight the importance of movement requirements and time constraints. Whereas reaches to stationary objects required a fast but self-determined movement, dynamic objects rolled rapidly toward a contact zone, providing an external cue for urgent reaches. The authors conclude that internally-regulated movements are more impaired in PD patients than externally-stimulated movements. Accordingly, PD patients showed similar eye and hand movements as controls during our track-intercept task which required urgent interceptive movements toward a designated strike zone. The task incorporated an external movement cue (the strike zone) and visual performance feedback—additional factors that might have facilitated preservation of function. Eye movements were also preserved in our baseline pursuit task, which required no urgency or deliberation similar to previous studies that tested simple ramp-pursuit tasks (Fukushima et al., 2013; 2015). These findings indicate that providing external stimulation— either through a task-evoked sense of urgency and temporal movement cues or through continuous stimulus presentation—is associated with preservation of eye and hand movements function in PD patients.

### Is sensorimotor prediction impaired in PD patients?

When interacting with moving objects, it is critical to accurately predict the sensory outcome of visual events (Fiehler et al., 2019). We tested participants in two tasks involving moving stimuli that required different levels of prediction. In the baseline pursuit task participants tracked a moving target that moved continuously and predictably. In the track-intercept task participants had to extrapolate the target’s trajectory after it had disappeared, requiring deliberate eye movements and interception at a predicted location. In both tasks, we found relative preservation of pursuit velocity and position error as well as preserved predictive ability to guide an interceptive hand movement.

By contrast, smooth pursuit had been shown to be impaired in task conditions that required integrating cue information or anticipation. When remembering the meaning of two consecutive cues, one direction cue and one go/no-go cue, PD patients tended to track the target using saccades rather than following it smoothly (Fukushima et al., 2013; 2015). Internally-generated or predictive movements were also impaired in studies using anticipatory pursuit in response to a target direction reversal (de Hemptinne et al., 2013) or target blanking (Helmchen et al., 2012), or when testing the accuracy of manually controlling a randomly moving target by using a joystick (Chen et al., 2016). These studies provide converging evidence that PD patients lose the ability to move in anticipation of a future visual event when tasks require concentration and effort but no implied urgency to move. In contrast, the combination of an externally-provided end location and a time-critical movement constraint (Majsak et al., 1998; 2008; Fooken & Spering, 2019; 2020) can facilitate the preservation of predictive abilities in PD patients.

### Brain networks underlying differential impairments in PD patients

Different levels of functional impairments in response to different types of visual stimulation have also been observed in healthy aging. For example, a study investigating motion perception in a large sample of healthy adults across the lifespan (Billino et al., 2008) found preserved ability to perceive complex motion patterns (biological motion and radial motion) as compared to simpler ones (translational motion). The authors speculate that motion stimuli with high ecological relevance (e.g., expanding radial flow might induce a fight or flight response) might be processed more efficiently, and potentially by a set of functional pathways that bypass primary visual cortex. Studies that found dissociations between motion perception and smooth pursuit eye movements have similarly argued that the pursuit system could be aided by a separate subcortical pathway that forms a direct connection from the retina to SC and brainstem (Spering & Carrasco, 2015).

Stimulus-dependent preservation and impairments of movements in PD is in accordance with the idea of different functional pathways. Dysfunction of the fronto-basal ganglia network might be linked to impaired inhibitory control of action planning and deliberation (Alexander & Cruther, 1990; Aron et al., 2007; Brown et al., 2004; Lalo et al., 2008; Mink, 1996; Wiecki & Frank, 2010). Preserved fast visuomotor responses, such as manual interceptions, and visually-guided eye movements might be associated with SC-brainstem loops (Corneil and Munoz, 2014) and the tecto-reticulo-spinal pathway (Gu et al., 2016). Preservation of oculomotor function in PD could also be mediated by a direct pathway, bypassing dopaminergic connections through the basal ganglia (Basso, Pokorny, & Liu, 2005) or a hyperdirect pathway linking cortical eye movement areas to the subthalamic nucleus of the basal ganglia, (Nambu et al., 2002; Sieger et al., 2013). The subthalamic nucleus is involved in pursuit and saccadic eye movement control and is a target area for deep brain stimulation in PD patients (FitzGerald & Antoniades, 2016; Lee et al., 2019).

Movement preservation and impairment in response to different types of stimuli and temporal task constraints might also be related task motivation. Previous research has linked bradykinesia in PD to a lack of movement motivation (Mazzoni et al., 2007). When patients were given feedback about their movement speed, they were able to point to a stationary target as fast and accurately as age-matched control. However, PD patients implicitly chose to move at a slower speed compared to controls and needed more repetitions to attain the desired number of valid (sufficiently fast) trials. The authors propose that impaired movement motivation is linked to dopaminergic projections from the midbrain to the striatum (Mazzoni et al., 2007; Niv et al., 2007; Schultz, 2007; Moustafa et al., 2008). Dopaminergic medication enhanced the ability of PD patients to anticipate error signals when continuously tracking an unpredictably moving visual target with a joystick (Chen et al., 2016), indicating that dopamine increases sensitivity to positive reinforcement learning processes (Frank et al., 2004). In our tasks, we did not find systematic effects of dopaminergic medication on any eye or hand movements. These findings are consistent with other studies showing comparable smooth pursuit eye movements in patients on and off medication (Fukushima et al., 2015; Ladda et al., 2008). A lack of medication effect on select oculomotor performance at an early stage in the disease might indicate that externally-stimulated movements (e.g., visually-guided eye movements) are less affected by a decrease of movement motivation.

## Conclusion

The present study provides evidence for stimulus- and task-dependent oculomotor deficits in PD patients. Systematic impairments of saccades to stationary targets at short latencies indicate impaired inhibitory oculomotor control in PD patients. In turn, the relative preservation of visually-guided smooth pursuit, motion prediction, and fast manual interception might be mediated by separate functional pathways rather than differences in movement motivation. Our findings can inform the development of tasks that are engaging and motivating for functional training in PD patients. Furthermore, we found evidence for adaptive mechanisms in the eye (decreased inhibition to compensate increased latency) and in the hand (decreased latency to compensate decreased velocity). Such long-term sensorimotor adaptation might be related to continuous reinforcement that patients receive during everyday life.

## Acknowledgements

This work was supported by a Deutsche Forschungsgemeinschaft (DFG) Research Fellowships to JF (grant FO 1347/1-1) and an NSERC Discovery Grant and Accelerator Supplement to MS. The authors thank members of the Spering lab for feedback on the manuscript.

